# Variant classification guidelines for animals to objectively evaluate genetic variant pathogenicity

**DOI:** 10.1101/2024.09.17.613537

**Authors:** Fréderique Boeykens, Marie Abitbol, Heidi Anderson, Iris Casselman, Caroline Dufaure de Citres, Jessica J. Hayward, Jens Häggström, Mark D. Kittleson, Elvio Lepri, Ingrid Ljungvall, Maria Longeri, Leslie A. Lyons, Åsa Ohlsson, Luc Peelman, Pascale Smets, Tommaso Vezzosi, Frank G. van Steenbeek, Bart J.G. Broeckx

**Author notes:** **Correspondence:** Bart J.G. Broeckx. These authors contributed equally to this work and share last authorship (Frank G. van Steenbeek, Bart J.G. Broeckx).

## Abstract

Assessing the pathogenicity of a disease-associated variant in animals accurately is vital, both on a population and individual scale. At the population level, breeding decisions based on invalid DNA tests can lead to the incorrect exclusion of animals and compromise the long-term health of a population, and at the level of the individual animal, lead to incorrect treatment and even life-ending decisions. Criteria to determine pathogenicity are not standardized, hence no guidelines for animal variants are available. Here, we developed and optimized the animal variant classification guidelines, based on those developed for humans by The American College of Medical Genetics and Genomics, and demonstrated a superior classification in animals. We described methods to develop datasets for benchmarking the criteria and identified the most optimal in silico variant effect predictor tools. As the reproducibility was high, we classified 72 known disease-associated variants in cats and 40 other disease-associated variants in eight additional species.

## 1 Introduction

The pace at which disease-associated variants in animals are discovered is increasing and is associated with technological advancement(1) (Fig. 1). The challenge of the interpretation of DNA variants and the translation to the clinic is well known in human medicine(2–5). The functional mechanisms of identified disease-associated DNA variants are often unclear and the associated diseases may have variable expression and incomplete penetrance, which leads to ambiguous interpretation of the pathogenicity of given variants. In animals, these challenges have also been recognized, nevertheless, no standardized evaluation protocol of variant pathogenicity has been developed(6–8). The potential consequences associated with misinterpretation of the importance of genetic variants however are far reaching(9–11). Ranging from incorrect treatment to even euthanasia, individual animals can suffer directly, however, the consequences can even negatively affect the entire population by impacting breeding decisions(9–11). As genetic diversity in several cat and dog breeds is low compared to the general human population, the exclusion of animals based on invalid associations can drive a further increase in other disease prevalences, and substantiate the concerns linked to animal welfare(12–18).

**Fig. 1.**
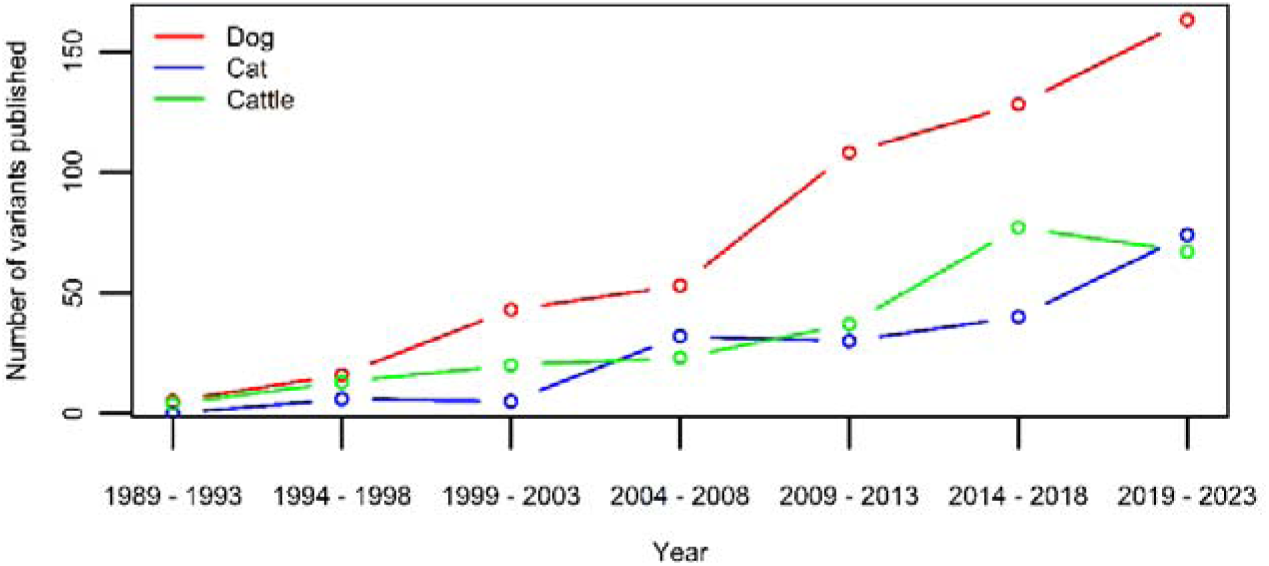
Number of new variants published per five year-period for the three species with currently over >100 published disease-causing variants in OMIA (1). While the extent is species-dependent, an increase can be seen in all three.

The American College of Medical Genetics and Genomics (ACMG) developed, and updates, widely used guidelines in human genetics to provide guidance when interpreting the potential pathogenicity of genetic variants(2,19–28). While these ACMG guidelines have been considered a few times more recently in animal genetics, their implementation is overall very limited, and concerns about the appropriateness of some criteria have led to the exclusion of certain criteria in some publications(29–32). More recently, applying the ACMG guidelines has been objectively shown to lead to misclassification of variants in cats(6). Currently, no guidelines have been established that assist the process of variant classification in animals, which implies that decisions on pathogenicity are not standardized and are based on individual investigator experience alone.

We present guidelines tailored for variant classification in domestic animals and demonstrate their superiority on animal variant classification relative to the ACMG guidelines. To benchmark these animal variant classification guidelines (AVCG), a reference dataset was created, with allele frequencies (AF) derived from a large population study. An evaluation of *in silico* variant effect predictor tools was performed. The reproducibility of labelling variants in a five-category classification system was examined and variants in nine species were evaluated.

## 2 Materials and methods

### 2.1 Ethics

Samples were non-invasive buccal swabs, collected by the owner or in veterinary practices, or whole blood on EDTA, collected by veterinary clinics, in accordance with international standards for animal care and research. In some cats, blood was obtained as part of routine clinical procedures for diagnostic purposes, at the request and with the consent of the owner. As these samples were from client-owned cats for which no harmful invasive procedures were performed, there was no animal experimentation according to the legal definition in Europe (Subject 5f of Article1, Chapter I of the Directive 2010/63/UE of the European Parliament and of the Council). All cat owners provided consent for the use of their cat’s DNA sample in scientific research. Written informed consent was obtained from the owners for the participation of their animals in this study.

### 2.2 Samples

The feline AF dataset was based on non-invasive cheek swab samples collected by cat owners, and either blood or cheek swab samples collected at certified veterinary clinics, for submission to commercial DNA testing. The samples submitted for MyCatDNA^TM^ / Optimal Selection^TM^ Feline (Wisdom Panel, Helsinki, Finland and Wisdom Panel, Vancouver, WA, USA, respectively) DNA genotyping based on a custom-designed Illumina® Infinium HD array between September 2016 and November 2023 consist of 30,577 individual cat samples, of these 11,036 samples (36%) have been previously published(33) and 19,541 samples are new entries. The samples submitted for Antagene DNA genotyping consist of 32,841 individual cat samples (Antagene, La Tour de Salvagny, France). The owners provided written consent for data use in research upon submission of samples for commercial genetic testing. All tests are routinely run and offered commercially.

The breed of a cat was reported by its owner typically with additional accompanying information confirming registration under The International Cat Association, Fédération Internationale Féline, Livre Officiel des Origines Félines, The Cat Fanciers’ Association, or World Cat Federation standards. Additional breeds not yet recognized by any major breed registry but with an established community of breed hobbyists were also considered breeds for the purposes of this study. A cautionary note specifying examples regarding this aspect was added (Suppl. Table S1). The non-pedigreed cat sample set consisted of mixed breeds, breed crosses, or random-bred cats.

### 2.3 The decision-making process to obtain variant classification guidelines

In agreement with recommendations assisting the development of guidelines, the following steps were taken(34,35). Prior to the actual development process, the need for the development of new guidance and evaluation and/or modification of existing guidelines for variant classification was decided on by the current multidisciplinary group. The starting point for this process were the original 2015 ACMG guidelines used in human medicine (Suppl. Table S2) (2). Throughout the manuscript, when a criterion from the original ACMG guidelines is mentioned, the criterion name will always be preceded by ACMG. If this is not the case, the criterion mentioned is part of the newly developed AVCG.

A multidisciplinary group was composed including veterinarians, geneticists from universities, and from commercial laboratories offering DNA-tests, and fell within the recommended six to fifteen members(35). The development process was divided into three phases: the pre-meeting preparatory phase (phase one), a group decision phase (phase two), and the optimization phase (phase three)(34,35). The first phase was an individual evaluation phase, whereas the focus group approach was used in phase two and three. In the first phase, each member of the multidisciplinary team independently provided remarks and voted (accept, revise, or remove) on each of the ACMG criteria, as well as had an opportunity to propose new criteria. The remarks and votes from phase one were shared anonymously in the decision phase (phase two). Decisions to retain, revise, or exclude ACMG criteria for AVCG were made by anonymous voting and were based on a two-thirds majority rule. Modifications were linked to the text of the criterion and/or the weight assigned to that criterion. Finally, in the optimization phase, individuals who worked with the guidelines could propose changes, which were again voted on anonymously by the entire group (phase three). All group meetings (phase two and phase three) were recorded.

While various classification development approaches exist, the focus group approach was chosen in phases two and three as the goal was to opt for a method that allowed 1) a group discussion to stimulate new ideas and insights (requirement one), and 2) a safe decision environment in which personal opinions can be reflected (requirement two)(36). While focus groups allow a thorough discussion and enrichment of ideas, potential drawbacks are the lack of confidentiality and power relations(36). To resolve that and fulfil requirement two, voting was anonymous in phases two and three and participants were invited to submit remarks prior to the decision phase, and these were also included anonymously in the presentations used in phase two. During the entire process, the stepwise development was tracked.

### 2.4 Obtaining a dataset of pathogenic and benign variants for benchmarking

To evaluate 1) the performance of both the ACMG and the AVCG, and, 2) the accuracy of *in silico* tools, datasets of “true” disease-causing variants and benign variants had to be created. A systematic review according to the PRISMA guidelines was conducted to evaluate the existing methodologies for selecting pathogenic and benign variants in studies involving these tools(37). The process is explained in Suppl. Data S1, graphically represented in Suppl. Figure S1 and methods are summarized in Table 1.

**Table 1.** Overview of variant selection methods used in benchmarking studies.

| Method | Comments | Suitable? |
| --- | --- | --- |
| **A. Pathological variants** |  |  |
| Database (e.g., Clinvar, Humsavar) | Usually based on a pathogenic/likely pathogenic label assigned with the ACMG classification. While the number of databases in veterinary medicine is limited, a database like OMIA is currently the closest proxy. | ✓ |
| Functional studies (e.g., RNAseq, enzyme assays) | Conducted when there is a focus on one specific phenotype and with limited number of variants linked to that phenotype. Taking the wide range of phenotypes and variants into account, this approach is not feasible. | ✗ |
| Literature review | Classification based on published literature. To decrease risk of misclassification, three independent, experienced reviewers, with only one reviewer per institution, reviewed the literature. | ✓ |
| **B. Benign variants** |  |  |
| Allele frequency | While the exact cut-offs used in human studies vary greatly (e.g., >1%, >40%, ...), the suitability of allelic frequency cut-offs in veterinary medicine has been heavily debated. Furthermore, first basing selection of genetic variants on allele frequency criteria and next using these allele frequency criteria for variant classification results in a circular reasoning that will bias the results and should thus be avoided. | ✗ |
| Database (e.g., Clinvar, Humsavar) | Often based on a benign/likely benign label assigned with the ACMG classification. No “benign/neutral” database is available in veterinary medicine. | ✗ |
| Functional studies (e.g., RNAseq, enzyme assays) | Conducted when there is focus on one specific phenotype and with limited number of variants linked to that phenotype. As several genes are expressed in various tissues and an individual variant can have an effect in various ways, the investigation of the potential absence of an effect is beyond the scope of most gene/disease investigations. | ✗ |
| Random set of variants | Taking into account that the ratio of disease-causing variants relative to the total number of variants in a variant database is extremely small, this strategy starts with random selection of variants from a variant database. Subsequently, to reduce the probability of erroneously including disease-associated variants further, we excluded variants linked with disease in other species, including humans. | ✓ |

Abbreviations: ACMG = American College of Medical Genetics and Genomics, OMIA = Online Mendelian Inheritance in Animals, ✓ = suitable, ✗ = not suitable.

Based on the summarized methods (Table 1), the following procedure was followed to collect a set of pathogenic variants. Every geneticist was asked to independently “provide at least five variants that he/she considers to be pathogenic without a doubt”. A list was compiled, removing all duplicates. Subsequently, these variants were manually checked and were excluded if they met any of the following exclusion criteria: somatic variants associated with cancer, variants (risk/protective) associated with complex traits, structural variants (defined as sequence variants >50bp in size(38)). Next, variants were removed if 1) the original paper could not be retrieved and/or 2) if there were errors in the paper and/or if 3) the paper was considered to be too old to make a proper judgment, and/or if 4) the variant caused a non-disease Mendelian phenotype.

To be retained in the final list, variants had to pass all the aforementioned exclusion criteria and consistent and independent labelling as “without a doubt pathogenic” by at least three experienced (defined as >10 years of experience in the field) geneticists. If a variant was put on the list independently by three geneticists during the collection process, the variant was considered pathogenic. If a variant was proposed by less than three geneticists, other geneticists than the ones that put it on the list, were asked to review the variant. If the variant was only reported once, two other geneticists were consulted, if it was reported twice, then only one other geneticist was consulted. Variants were allocated to reviewing geneticists randomly and during the entire process (submission for the initial list and reviewing of the remainder) no instructions were given on the methodology, i.e., every geneticist did this at their own discretion. The concordance of the independent evaluations was tracked.

This dataset was completed with AF data as this is an important part of the evaluation process in the ACMG guidelines. To provide a standardized and as complete dataset as possible, allelic frequency data from two commercial laboratories (Antagene and Wisdom Panel) were added for every variant that was routinely tested (for details: see “2.2 Samples”).

Similarly, based on the results of the systematic review according to the PRISMA guidelines (strategy detailed in Suppl. Data S1, graphically represented in Suppl. Figure S1 and methods summarized in Table 1), the following selection strategy was used to retrieve variants that were most likely benign(37). First, a random selection of a variant with a certain effect was done from BioMart(39). The following settings were used: 1) Database: Ensembl Variation 111, Cat short variants (SNPs and indels excluding flagged variants) (Felis_catus_9.0); 2) Filters: Region: none; General variant filters: none; Gene associated variant filters: “missense_variant”, “frameshift_variant”, “stop gained”, 3) Attributes: variant associated information: variant name, variant source, chromosome/scaffold name, chromosome/scaffold position start (bp), chromosome/scaffold position end (bp) (default settings). The results were exported and a random selection was performed in R version 4.3.2 using a custom script. Second, the gene in which the selected variant resides was checked for a reported association with disease in Online Mendelian Inheritance in Animals (OMIA)(1). Third, a homology-based search through ConVarT was used to identify whether a variant in humans was reported to be pathogenic or likely pathogenic according to ClinVar(40). Only when OMIA yielded no result and the variant was not classified pathogenic or likely pathogenic in ClinVar, the variant was retained in the list. This process was continued until a number equal to the number of pathogenic (missense/nonsense/frameshift) variants was achieved. This list is available in Suppl. Table S3. For splice sites, this approach was not used. As there is no functional proof that a random splice variant from the database will actually result in altered splicing, neither that the absence of such a variant will not, it would not be possible to check whether a tool correctly predicts the effect. However, as the splice site variants from the pathogenic dataset contained both the creation and removal of donor as well as acceptor sites and functional proof for the effect (i.e., cDNA/mRNA sequence evaluation) for seven out of eight variants was provided, these were used to assess the predictions of the tools.

The pathogenic and benign dataset were restricted to one species to avoid potential bias due to species differences; species transferability of AVCG was evaluated in a subsequent dataset (see “2.8. Evaluation of the performance of variant classification guidelines: cross-species classification”). From the nine species mentioned individually on OMIA, the species selected had to have a sufficiently large number of disease-associated variants published (>100) and to ensure maximum confidence in the dataset, the geneticists involved in the variant evaluation, had to have ample experience with that species.

### 2.5 Selection, analysis, and optimization of the *in silico* variant effect predictor tools

A systematic review according to the PRISMA guidelines was conducted to obtain a list of *in silico* tools(37). The procedure is described in Suppl. Data S2 and graphically represented in Suppl. Figure S2. After the initial list was compiled, all tools were subsequently evaluated for 1) easy accessibility, defined as whether there is an online interface available (i.e., no download is necessary to use the tool), and 2) whether non-model species were supported. The remaining tools were all included in the benchmark analyses.

For missense, nonsense and frameshift variants, to evaluate performance, a balanced design was used, i.e., an equal number of pathogenic and benign variants was included during benchmarking. For each tool, the number of variants that gave a result, as well as the accuracy, sensitivity, and specificity were calculated. The default settings were used. While most tools dichotomize classification, PANTHER and PolyPhen2 have subcategories (probably/possibly benign or damaging, respectively), which were collapsed into pathogenic or benign(41–43). For splice sites, reporting is slightly different: the output of the tools is not pathogenic or benign, but whether a donor or acceptor splice site is created or not. As such, the overall accuracy is reported.

Whenever >1 tool was available to assess a specific category of variants, the most optimal combination of two tools was identified to adhere to the *in silico* criterion as mentioned in ACMG and the AVCG. For missense, nonsense and frameshift variants, most optimal was defined as the combination of tools that led to the least consistent misclassification, i.e., the least false positives (variants from the benign list that were allocated a “pathogenic” label) and false negatives (variants from the pathogenic list that were allocated a “benign” label) relative to the total number of consistent classifications. If there was a tie, the combination that classified most variants correctly, was preferred. For splice sites, the best combination of tools was defined as the two tools that together consistently predicted most often correctly the creation/removal of donor/acceptor sites, respectively.

### 2.6 Evaluation of the performance of variant classification guidelines: classification of pathogenic variants

The variants in the aforementioned “truth” pathogenic variant list were classified twice: once with the traditional ACMG guidelines and once with the newly developed AVCG guidelines(2). To avoid the tendency to look for additional criteria supporting pathogenic classification if the threshold would not be met, the reviewers were only asked to check the criteria and not to calculate the final label. No variants were reviewed by the same investigator who made the original variant discovery. Furthermore, the work was divided, where each time two geneticists reviewed the *in silico* tools, two others reviewed the newly added criteria and the final two reviewed the remaining criteria. Label assignment was finally done by a different geneticist who adhered to the decision table exactly.

Practically this entails that for each variant, all the evidence of the various criteria that were fulfilled, was weighted and counted to determine a classification. While the decision-making process changed slightly from ACMG to AVCG, a five-category-based classification remained (Table 2)(2). The categories/labels are: pathogenic (P), likely pathogenic (LP), variant of unknown significance (VUS), likely benign (LB) and benign (B). Each category has consequences for clinical decision making and for whether a variant should be included in breeding and/or screening programs. As primarily differences between P/LP and the other three categories might lead to differences in medical management and/or breeding strategies, these are specifically mentioned in subsequent analyses, aside from general classification overviews(3).

**Table 2.** Description and consequences of each variant pathogenicity classification in the five-category system.

| Label | Consequence |
| --- | --- |
| Pathogenic (P) | a healthcare provider can use molecular testing information in clinical decision-making, for breeding programs and/or screening. |
| Likely pathogenic (LP) | a health-care provider can use the molecular testing information in clinical decision-making when combined with other evidence of the disease in question, for breeding programs and/or screening. |
| Variant of uncertain significance (VUS) | not to be used in clinical decision-making, for breeding programs or screening. Efforts to resolve the classification of the variant as pathogenic or benign should be undertaken. |
| Likely benign (LB) | a healthcare provider can conclude that it is not the cause of the patient's disorder when combined with other information. It should not be used for breeding programs and/or screening. |
| Benign (B) | a healthcare provider can conclude that it is not the cause of the patient's disorder. It should not be used for breeding programs and/or screening. |

Each category is linked to specific recommendations in terms of clinical decision making, inclusion in breeding and screening programs. Between P and LP, mainly the strength of the evidence varies, while the practical differences are limited. Similarly, the practical differences between VUS, LB, B also have limited consequences. The most significant health and breeding decisions would occur when a variant switches between the pathogenic (P/LP) and the benign/uncertain classifications (VUS/LB/B), which can happen when new information becomes available or when label assignment differs between evaluators, hence the importance of a high inter-evaluator agreement on classification.

### 2.7 Evaluation of the performance of variant classification guidelines: inter-evaluator agreement of classification

Five evaluators independently assessed a random subset of an overlapping set of variants with the newly developed guidelines. The goal was to provide a set that encompassed variants with various effects (missense/ nonsense/ frameshift/ splice sites) and that vary from easy to more difficult to diagnose (variants belonging to the set of pathogenic variants (n=6), variants that were submitted for the set of pathogenic variants but that were not consistently considered pathogenic by all three evaluators (n=5) and a set of variants for which the ACMG criteria turned out to be unsuitable, resulting in a wide range of classifications back then (n=6))(6). To ensure that classification differences were linked to differences in the interpretation of the guidelines rather than differences in terms of access to data linked to the variant, all evaluators based their classification on the same papers. In total, 17 variants were evaluated by three geneticists independently.

To evaluate the inter-evaluator agreement, all pairwise combinations were checked. Classifications that might lead to medical management differences (i.e., P/LP versus B/LB/VUS) and disagreements less likely to affect clinical decision-making (P versus LP; B versus LB; VUS versus LB/B), were evaluated.

### 2.8 Evaluation of the performance of variant classification guidelines: cross-species classification

While these guidelines were initially tested on a set of feline variants, the goal was to develop a set of variant classification guidelines capable of classifying variants across a wide range of domestic species. Aside from the cat, the applicability of AVCG across species was evaluated for all eight other species mentioned in OMIA(1). Practically, for every additional species, five variants were evaluated, if available, and the evaluator was asked to answer two questions: 1) whether there was an incompatibility of any of the criteria in that specific species in which that variant was evaluated and 2) whether they encountered difficulties not seen in the cat. To avoid any bias, no evaluator was allowed to check a variant published by his/her own group.

## 3 Results

### 3.1 The development of guidelines for variant classification

#### 3.1.1 Defining the scope

The scope was defined as the development of guidelines that are used by the community for Mendelian disorders in animals to classify variants in categories based on standardized criteria. Somatic variants associated with cancer, variants (risk/protective) associated with complex traits, and structural variants (defined as a variant larger than 50bp(38)) were considered outside the scope of these guidelines.

#### 3.1.2 Evaluation of the ACMG criteria

The development of the AVCG was conducted in three steps. After the preparatory phase, seven ACMG criteria were removed, while six were altered in phase two. In phase three, one additional criterion was altered. Overall, of the 28 initial criteria from the original ACMG guidelines published in 2015, half were removed or altered(2). The final set of criteria can be found in Table 3, clarifications for several criteria can be found in Suppl. Data S3.

**Table 3.** Criteria to support classification of pathogenic and benign variants in animals.

| Name | Criterion |
| --- | --- |
| PVS1 | Null variant (nonsense, frameshift, canonical $\pm 1$ or 2 splice-sites, initiation codon, single or multi-exon deletion) in a gene where LOF is a known mechanism of disease in the same or another species, if functionality of the gene is expected to be similar across species. |
| PS1 | Same amino acid change as a previously established pathogenic variant regardless of nucleotide change. |
| PS2 | *de novo* in a patient with the disease and unaffected parental samples tested negative. |
| PS3 | Well-established *in vitro* or *in vivo* functional studies supportive of a damaging effect on the gene or gene product. |
| PS4 | The prevalence of the variant in affected individuals is significantly increased compared with the prevalence in controls. |
| PS5 | Cosegregation with disease in multiple affected family members in a gene definitively known to cause the disease. |
| PM1 | Located in a mutational hot-spot and/or critical and well-established functional domain (e.g., active site of an enzyme) without benign variation across breeds and/or species. |
| PM2 | Novel missense change at an amino acid residue where a different missense change has been determined to be pathogenic in other individuals. |
| PM3 | For recessive disorders, detected in *trans* with a pathogenic variant. |
| PM4 | Protein length changes as a result of in-frame deletions/insertions in a non-repetitive region or stop-loss variants. |
| PP1 | Cross-species alignment shows the variant is conserved and other information across species (e.g., ClinVar data) states the variant is pathogenic. |
| PP2 | Missense variant in a gene that has a low rate of benign missense variation and in which missense variants are a common mechanism of disease. |
| PP3 | All computational evidence supports a deleterious effect on the gene or gene product (conservation, evolutionary, splicing impact, etc.). |
| PP4 | Patient's phenotype or family history is highly specific for a disease with a single genetic etiology. |
| BS1 | Lack of segregation in affected members of a family. |
| BS2 | Observed in a healthy adult individual for a recessive (homozygous), dominant (heterozygous), or X-linked (hemizygous) disorder, with full penetrance expected at an early age. |
| BS3 | Well-established *in vitro* or *in vivo* functional studies show **no** damaging effect on protein function or splicing. |
| BP1 | Cross-species alignment shows the variant is not conserved and other information across species (e.g., ClinVar data) states the variant is benign. |
| BP2 | Observed in *trans* with a pathogenic variant for a fully penetrant dominant gene/disorder or observed in *cis* with a pathogenic variant in any inheritance pattern. |
| BP3 | In-frame deletions/insertions in a repetitive region without a known function. |
| BP4 | All computational evidence supports a benign effect on the gene or gene product (conservation, evolutionary, splicing impact, etc.). |
| BP5 | Variant found in a case with an alternate molecular basis for disease. |
| BP6 | A synonymous (silent) variant for which splicing prediction algorithms predict no impact to the splice consensus sequence nor the creation of a new splice site AND the nucleotide is not highly conserved. |

The name of each criterion designates whether it supports pathogenic or benign classification of a variant (P or B, respectively), followed by the weight of the support (VS = very strong; S = strong; M = moderate; P = supportive, respectively) and a number that represents the criterion. Explanations per criterion are provided in Suppl. Data S3. Abbreviations: LOF = loss-of-function

An overview of the removed criteria is provided in Suppl. Table S2. From the seven criteria that were removed, three were associated with AF (ACMG PM2, ACMG BA1, ACMG BS1; Suppl. Table S2). The rationale for removal of these three criteria was based on a combination of observations. First, the large variant databases used by human geneticists, seldom exist in other species, hence the data will often not be available. Secondly, animal geneticists must consider breed population structures and dynamics. Breeds are not always consistently defined across breed registries, and their populations experience the effect of (a combination of) breeding practices like population bottlenecks, popular sire effects, and inbreeding, which all influence AF. The human criteria have not been designed for those situations. Thirdly, while methods have been developed to calculate AF cutoffs, they rely on estimates of prevalence and penetrance, which can lead to debate and different results(44,45). This causes undesired subjectivity in the criteria. While the first reason will become less important when more data is generated, the other two could currently not be resolved, hence the criteria were removed. Two criteria linked to the authority of a reputable source (e.g., investigators from an experienced laboratory make a statement about a variant without showing the data, ACMG PP5 and ACMG BP6) have also been removed, due to the absence of reviewable evidence supporting the claim. Information must be available to allow independent checks. One criterion (ACMG PM6) was removed as it became redundant due to the rewording of another criterion. The final one (ACMG BP1) was deemed too restrictive without sufficient evidence.

#### 3.1.3 Inclusion of new criteria

Two new criteria were proposed in phase one, their concept was accepted in phase two and their text was optimized and assigned a supporting weight in phase three. These two criteria are essentially the pathogenic and benign version of an approach integrating conservation and clinical data from other species (Table 3, criteria PP1/BP1). As such, the newly developed guidelines (Table 3) contain 23 criteria, of which 14 are linked to pathogenicity and nine support benign classification, and these are combined to assign pathogenicity labels as detailed in Table 4. This also implies that there are 16 changes in AVCG versus the ACMG guidelines (i.e., 14 criteria removed or altered, two new criteria added).

**Table 4.**
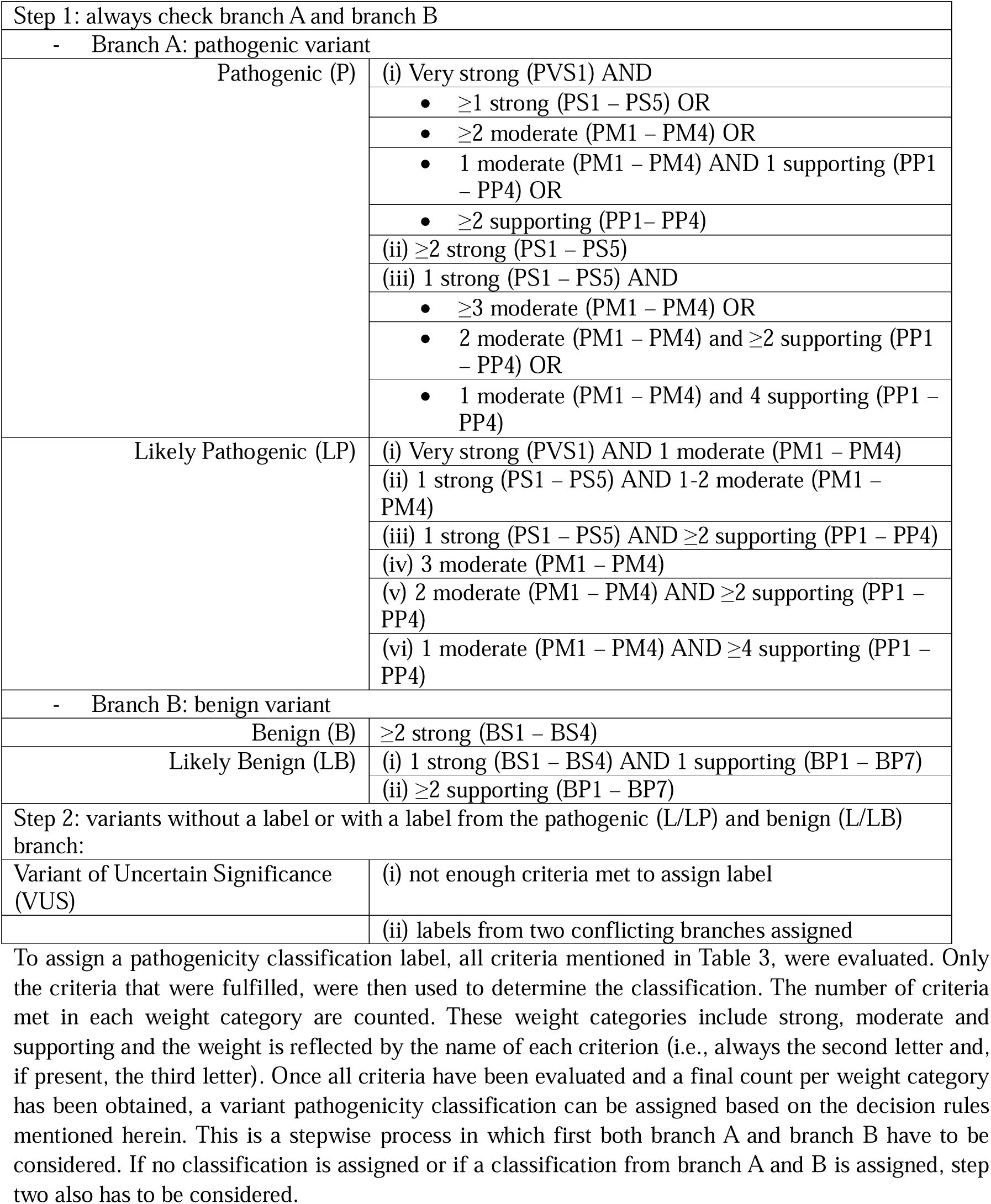
Decision rules to assign a genetic variant pathogenicity label.

### 3.2 Obtaining datasets for benchmarking

#### 3.2.1 A feline pathogenic variant “truth” set

As the cat is one of the top three domestic species for which disease-associated variants are published and several geneticists are dedicated feline experts, the cat was the species of choice for this analysis(1). The systematic review conducted to identify criteria used to search for candidate pathogenic variants in variant effect predictor benchmark studies (Suppl. Figure S1 for the PRISMA flow diagram), led to a set of 61 descriptions on how to obtain pathogenic variants (Suppl. Table S4). Summarizing the approaches led to three selection methods (Table 1), of which two were found suitable and were combined. To increase the stringency further, it was decided that independent literature reviews had to be conducted by at least three experienced (>10 years of experience) geneticists. Ultimately, 53 feline variants, encompassing a variety of mechanisms, inheritance patterns, and phene classes were considered pathogenic “without a doubt” by at least three geneticists and were used as a “truth set”(46,47). The characteristics of this pathogenic dataset are summarized in Table 5 and a detailed description per variant is provided in Suppl. Table S5.

**Table 5.**
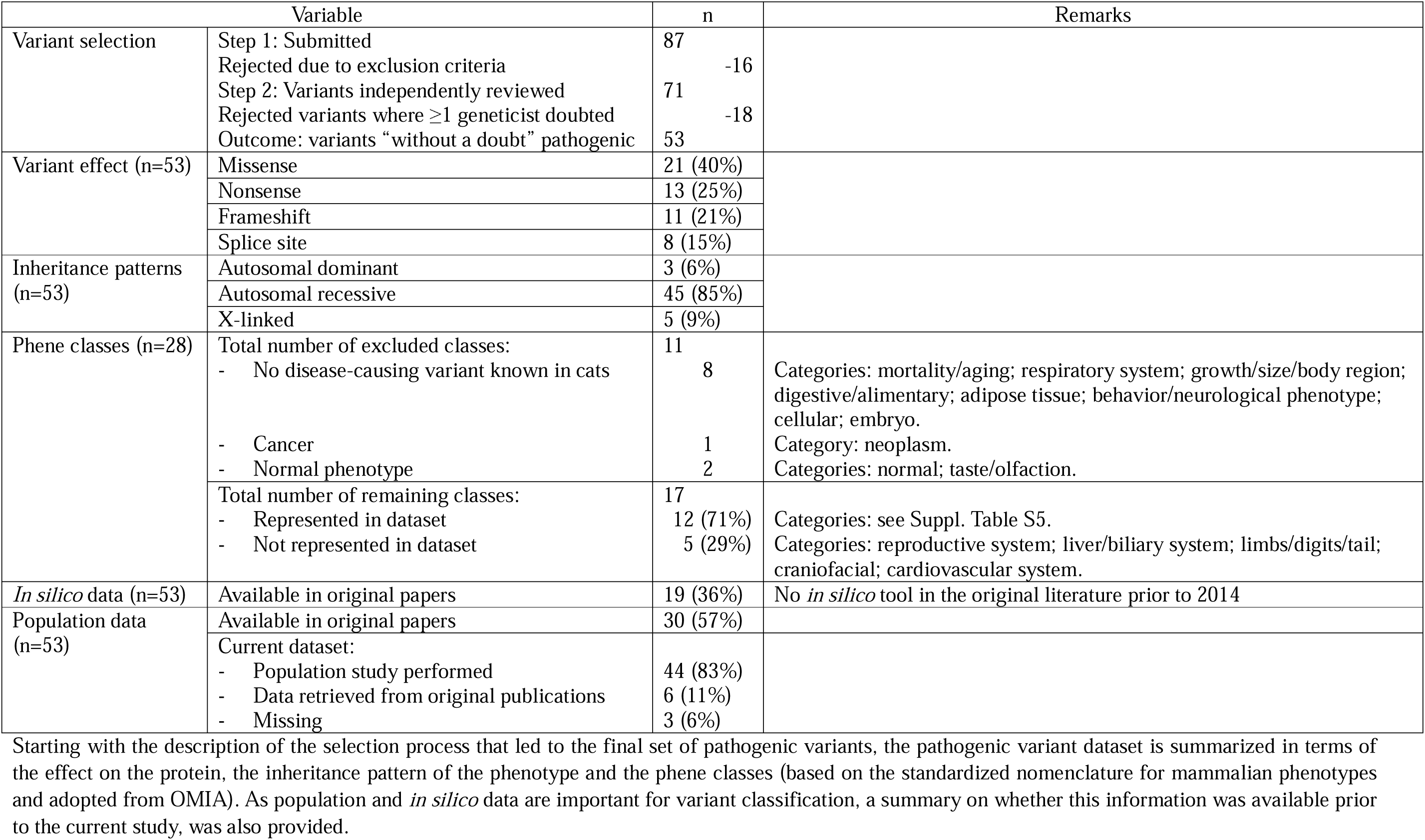
Characteristics of the feline pathogenic variant dataset.

Importantly for classification purposes, but also part of our recommendations on how variants should be reported (Suppl. Data S4), population data could be collected for 44 of the 53 variants, with sample sizes ranging from 994 to 48,949 (median: 31,559 samples), tripling the sample size from the largest feline population study so far(33). The AFs retrieved for these 44 variants ranged from 0% (i.e., not found in the population, 30/44 variants) to >5% (the variant responsible for factor XI deficiency) (Suppl. Table S5)(48). Seven variants were reported for the first time in at least one new breed (Suppl. Table S1)(49–53). Of the remaining nine variants, AFs for six of them were found in the original publication, leading to a total of 50 out of 53 variants with AF data (median sample size: 125, range: 28 - 597 samples). The complete list of variants and their overall and breed-specific AFs are provided in Suppl. Table S5 and S1, respectively.

#### 3.2.2 Development of a benign variant dataset

For several categories of variants (i.e., missense, frameshift, and nonsense variants, but not for splice site variants), *in silico* tools aim to predict whether they are disease-causing or not. To allow benchmarking of the performance of those tools, the aforementioned systematic review was conducted to identify criteria used to select candidate-benign variants (also described as “neutral”, “nondamaging” or “tolerated” in literature) for benchmark studies (Suppl. Figure S1 for the PRISMA flow diagram)(54). A set of 61 descriptions was obtained describing how to obtain benign variants (Suppl. Table S4). These descriptions could be summarized in four selection methods (Table 1) but only one method was found suitable. To obtain a balanced pathogenic-benign benchmark dataset, the selection process was continued until the number of feline benign variants matched the number of feline pathogenic missense, frameshift, and nonsense variants. The benign dataset can be found in Suppl. Table S3.

### 3.3 Analysis of *in silico* variant effect predictors

The systematic review returned 78 publications, identifying altogether 128 variant effect predictor tools (Suppl. Table S6). From this initial list of 128 tools, 114 were excluded because they did not allow analysis of genetic variants of cats and/or did not have a working online tool, leading to 14 tools that were ultimately retained. Of these, nine can be used solely to evaluate missense variants(55–63), one for nonsense and missense variants(55), one for nonsense variants and frameshifts(64), and three for splice sites(65–67), respectively. An overview of their performance (i.e., accuracy, sensitivity and specificity) and technical summary are provided in Table 6.

**Table 6.**
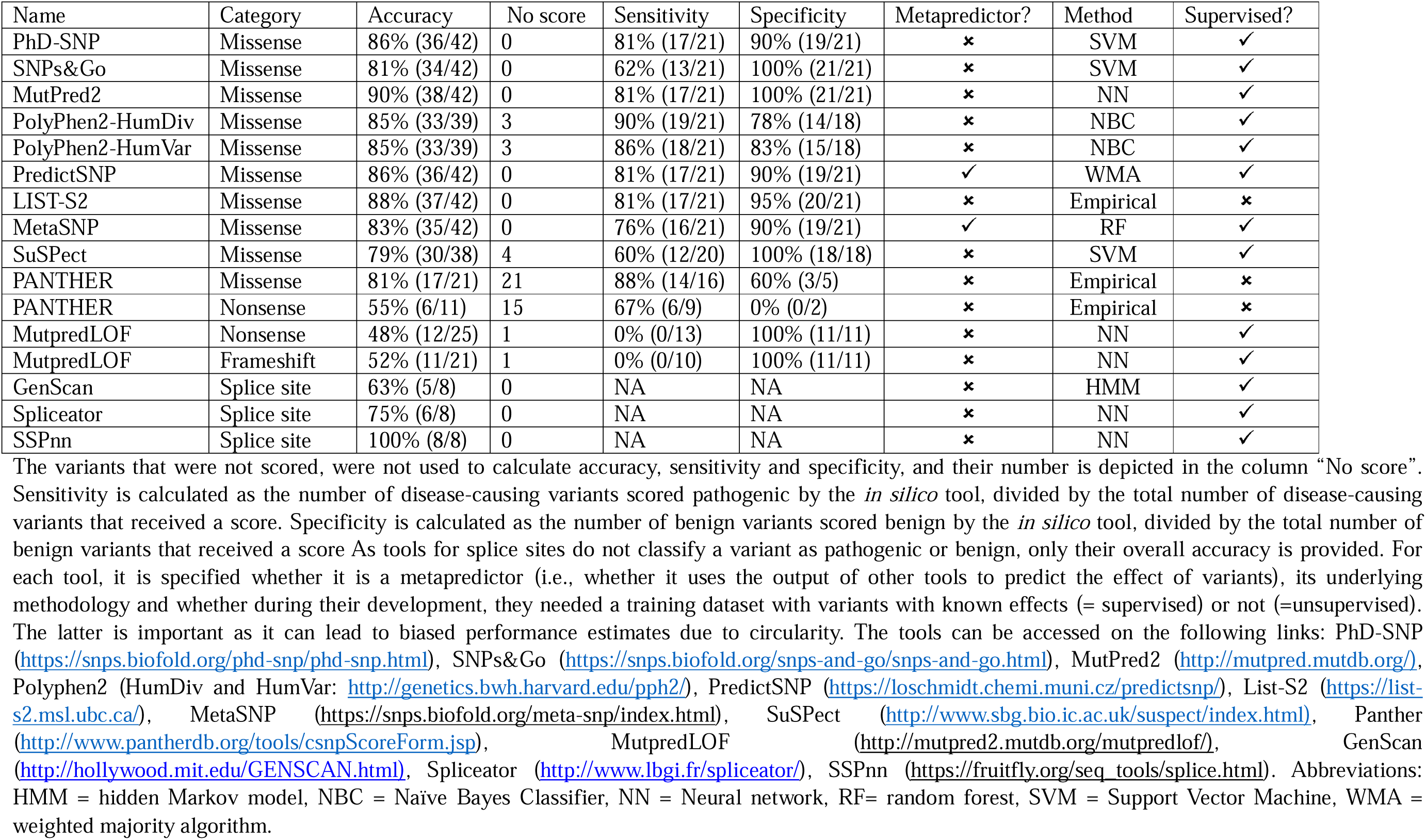
Overview of i*n silico* tools.

| Name | Category | Accuracy | No score | Sensitivity | Specificity | Metapredictor? | Method | Supervised? |
| --- | --- | --- | --- | --- | --- | --- | --- | --- |
| PhD-SNP | Missense | 86% (36/42) | 0 | 81% (17/21) | 90% (19/21) | ✖ | SVM | ✓ |
| SNPs&Go | Missense | 81% (34/42) | 0 | 62% (13/21) | 100% (21/21) | ✖ | SVM | ✓ |
| MutPred2 | Missense | 90% (38/42) | 0 | 81% (17/21) | 100% (21/21) | ✖ | NN | ✓ |
| PolyPhen2-HumDiv | Missense | 85% (33/39) | 3 | 90% (19/21) | 78% (14/18) | ✖ | NBC | ✓ |
| PolyPhen2-HumVar | Missense | 85% (33/39) | 3 | 86% (18/21) | 83% (15/18) | ✖ | NBC | ✓ |
| PredictSNP | Missense | 86% (36/42) | 0 | 81% (17/21) | 90% (19/21) | ✓ | WMA | ✓ |
| LIST-S2 | Missense | 88% (37/42) | 0 | 81% (17/21) | 95% (20/21) | ✖ | Empirical | ✖ |
| MetaSNP | Missense | 83% (35/42) | 0 | 76% (16/21) | 90% (19/21) | ✓ | RF | ✓ |
| SuSPect | Missense | 79% (30/38) | 4 | 60% (12/20) | 100% (18/18) | ✖ | SVM | ✓ |
| PANTHER | Missense | 81% (17/21) | 21 | 88% (14/16) | 60% (3/5) | ✖ | Empirical | ✖ |
| PANTHER | Nonsense | 55% (6/11) | 15 | 67% (6/9) | 0% (0/2) | ✖ | Empirical | ✖ |
| MutpredLOF | Nonsense | 48% (12/25) | 1 | 0% (0/13) | 100% (11/11) | ✖ | NN | ✓ |
| MutpredLOF | Frameshift | 52% (11/21) | 1 | 0% (0/10) | 100% (11/11) | ✖ | NN | ✓ |
| GenScan | Splice site | 63% (5/8) | 0 | NA | NA | ✖ | HMM | ✓ |
| Spliceator | Splice site | 75% (6/8) | 0 | NA | NA | ✖ | NN | ✓ |
| SSPnn | Splice site | 100% (8/8) | 0 | NA | NA | ✖ | NN | ✓ |

For 42 missense variants, from the ten remaining tools, six scored all 42 variants and the remaining four a subset. Most variants that were not scored were benign (25 times a benign variant versus six times a pathogenic variant). While the overall accuracy ranged between 79% and 90%, the best scores were for MutPred2 (90%), and List-S2 (88%). Two tools were available for analysis of 26 nonsense variants, PANTHER and MutpredLOF. MutpredLOF did not score one variant and PANTHER did not score 15 out of 26 variants. Again, more benign variants were not scored (12 times a benign variant versus four times a pathogenic variant). The overall accuracy ranged between 48 and 55%. For the 22 frameshift variants, only MutpredLOF was available. This tool scored all except one benign variant and the accuracy was 52%. For the eight splice site variants, three tools were available, with accuracies ranging between 63% and 100%.

As the *in silico* criteria used by ACMG and AVCG mention the combined use of several tools, if possible, each pairwise comparison was also assessed. This step was restricted to tools that assessed all variants in their category for two reasons: 1) to achieve general applicability, more versatile tools that score more variants can be more widely used, and 2) to avoid biased accuracy estimates because the omitted variants might be more difficult or easier to score.

Practically, if the results of two *in silico* tools are consistent, PP3 or BP4 are fulfilled. Otherwise, the result is not considered. Consistency does however not necessarily imply accuracy: two tools can be consistently wrong. As a wrongly fulfilled criterion can influence the final label and thus clinical decision-making and an inconsistent will not count in either direction, the “best” combination of *in silico* tools was defined as the combination that had the highest number of consistent correct classifications relative to the sum of consistent correct and consistent incorrect classifications. An overview of the combinations and their accuracy can be found in Suppl. Table S7. For missense variants, of 15 possible combinations, the combination that scored best was the combination of MutPred2 and List-S2 (34 out of 35 correctly classified). For nonsense and frameshift variants, this comparison was impossible as no tools scored all variants or only one tool was available, respectively. For splice site variants, the best combination was SSPnn and Spliceator.

### 3.4 Performance of the variant classification guidelines

#### 3.4.1 Classification of pathogenic variants

The 53 pathogenic variants were all classified with the ACMG criteria and AVCG. Overall, with the ACMG guidelines, 38 were classified as P (72%), six as LP (11%), four as VUS (8%), and five as B (9%), which implies that 83% of the variants were correctly classified. Overall, AVCG classified 42 variants as P (79%), seven as LP (13%), and the remaining four as VUS (8%); 92% of the variants were thus correctly classified, i.e., misclassification halved. The final classification for every variant is denoted in Suppl. Table S5.

As several variants were classified differently, the changed and/or removed criteria alter the classification. To investigate the differences, an in-depth evaluation was performed. Six variants were classified differently (Table 7)(48–50,52,68,69). For each of these, ACMG classified at least one category more in the direction of B. Five of these classification differences might affect clinical decision making and screening/breeding programs. An assessment of the differences demonstrated that this was linked to the removed AF criteria (ACMG BA1, five times negative effect), the newly added PP1 (two times positive effect), and the altered PM2 and PS5 (one and three times positive effect, respectively). The positive effects with AVCG were mainly linked to the specific focus on information available from other species. In short, four criteria resulted in a different classification and overall, the changes made during the development of AVCG, led to improved results.

**Table 7.**
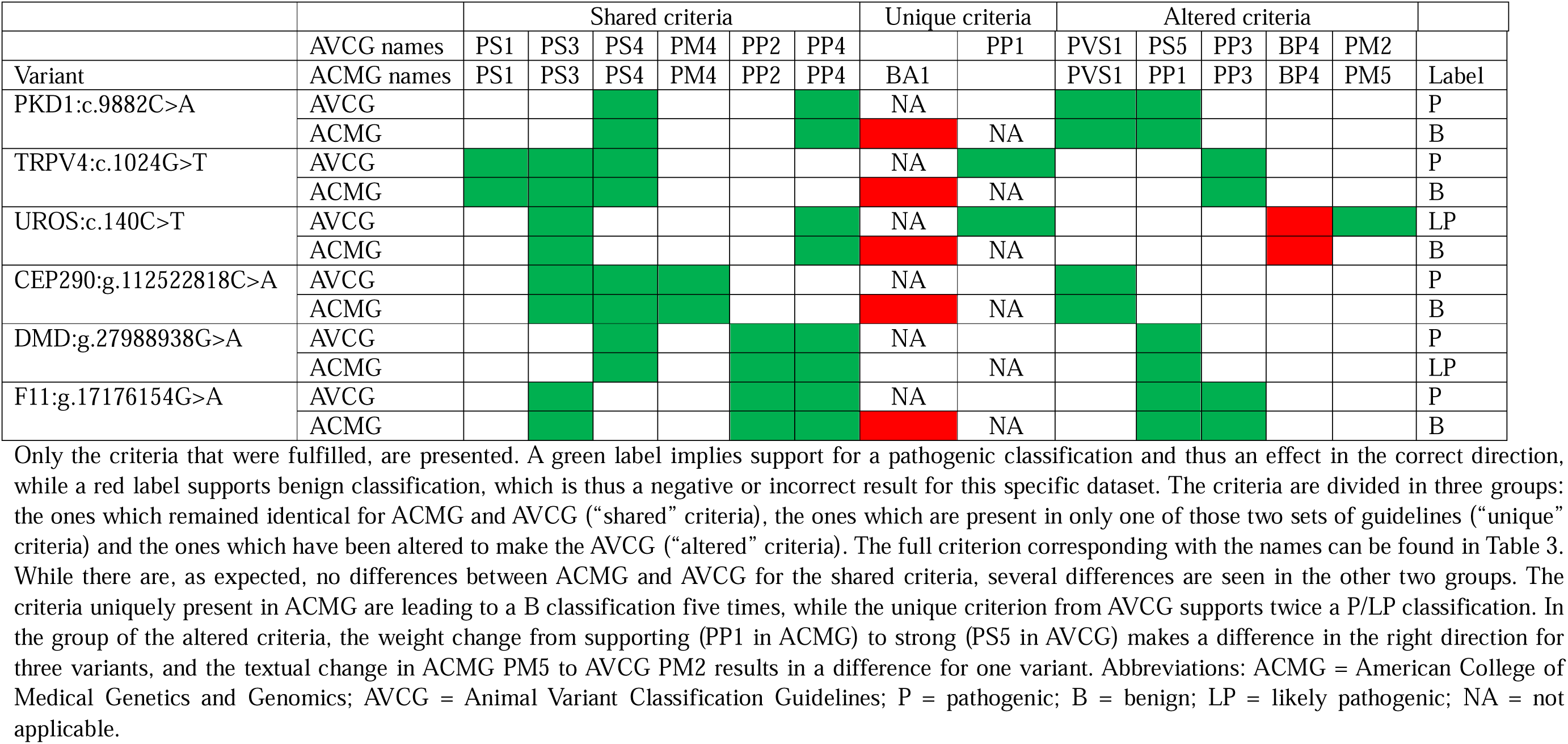
Variants classified differently with ACMG and AVCG.

An evaluation of the incorrectly classified variants (i.e., that did not get the label LP/ P) led to a total of nine variants that need to be scrutinized (Table 7-8). From these nine variants, five are uniquely misclassified with the ACMG guidelines (Table 7), while the other four were consistently misclassified as VUS (Table 8)(48–50,52,68,70–73). All five variants that were uniquely misclassified by ACMG were all directly caused by the AF criterion ACMG BA1 (Suppl. Table S2) which is removed in AVCG.

**Table 8.**
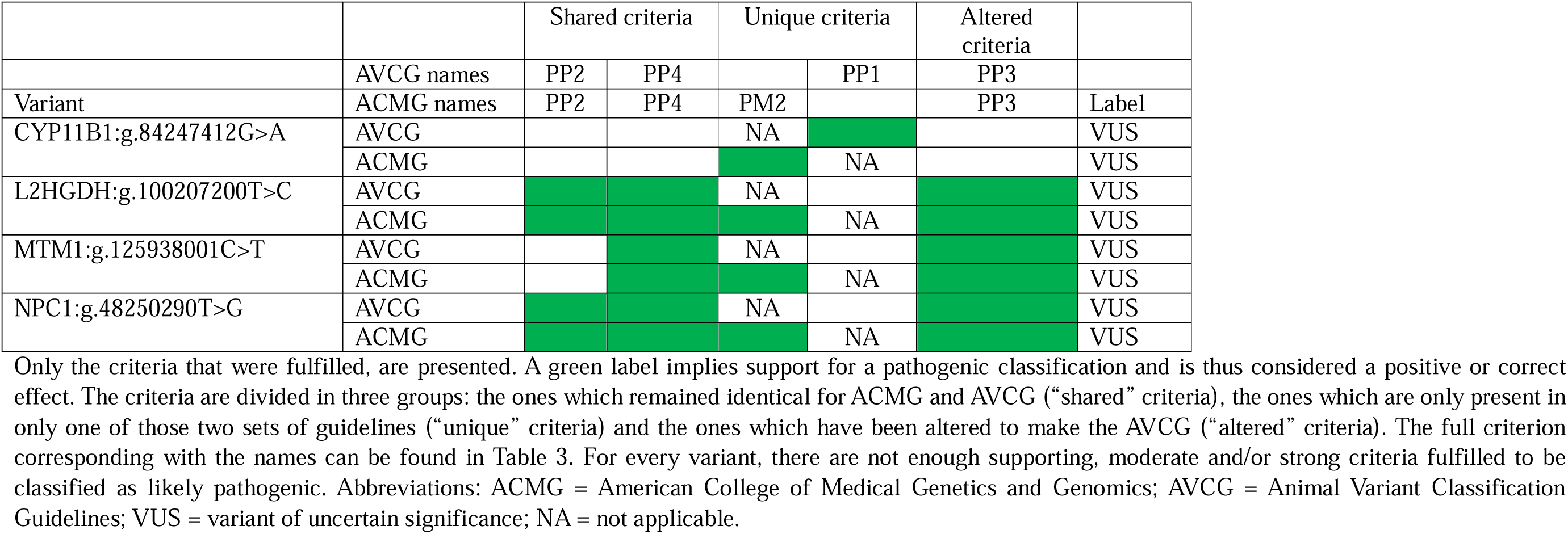
Variants consistently misclassified with ACMG and AVCG.

An analysis of the four VUS misclassified by ACMG and AVCG indicates that all four are missense variants with some criteria supporting pathogenic classification, however, not enough to be considered LP (Table 8). Another similarity is that they were all present in one unique case, i.e., there were no additional affected individuals or family members. As 15 other variants with n=1 were correctly classified (Suppl. Table S5), a detailed analysis was conducted to identify what sets them apart. What differs for the majority is their effect at the protein level: in 10/15, the variant is of a type invoking PVS1, a very strong criterion that only requires limited additional criteria to result in an LP/P classification. However, this still implies that five missense variants with n=1 were correctly classified (i.e., all as LP). For these five, this LP classification was due to an additional strong criterion (i.e., functional studies (PS3, n=2), an identical or other amino-acid change as a previously established pathogenic change (PS1, n=1; PM2, n=1, respectively)) or a combination of a sufficient number of moderate and supporting criteria being fulfilled.

#### 3.4.2 Inter-evaluator agreement of classification

Aside from correct classification, guidelines ideally also result in a consistent classification when considered by different evaluators. To assess this, a new dataset (Suppl. Table S8) was created, consisting of three groups of variants with different challenges in classification. Six variants belonged to the pathogenic reference dataset, six variants were included because ACMG classification in an earlier study(6) led to wide classification differences depending on which criteria were used, and for the remaining variants, there was a debate to whether these should be included in the pathogenic “truth” set. This dataset contains 17 variants and they were assessed with the AVCG by three individual evaluators, leading to a total of 51 classifications. These classifications were compared pairwise to evaluate differences.

Overall, 39 of 51 classifications (76%) matched. While there were a total of 12 of 51 disagreements on classification (24%), for only 2/51 (4%), this potentially might influence clinical decision-making (i.e., these were differences between P and VUS). For that specific variant, this was linked to a different evaluation of criterion PS3, i.e. whether there was sufficient *in vitro* or *in vivo* functional data to support a damaging effect. The remaining differences were linked to the strength of pathogenicity (8/51 (16%) P versus LP) or VUS versus benign classification (2/51, 4%).

Among the three groups of variants included in this reproducibility evaluation, clear differences in agreement were noticed (Suppl. Table S8). For the variants that passed the inclusion criteria for the pathogenic reference data set, the agreement was strongest (16/18 (89%)) and there were no misclassifications that might potentially lead to differences in medical management/ breeding/screening programs because they ranged between P and LP. Among the variants that were included because ACMG classification was difficult, the agreement was in the same range (14/18 (78%)) with also no misclassifications affecting clinical management. Among the variants that were included because there was a disagreement among evaluators whether they should be included in the pathogenic dataset, the agreement was lower (9/15 (60%)) and 2/15 differences (13%) might lead to altered clinical approaches as they ranged between VUS and P. For these five variants, this is still an increase in agreement compared to the 5/15 (33%) agreement based on subjective decision-making on pathogenicity prior to the development of the guidelines.

#### 3.4.3 Across-species classification of variants

As the AVCG worked well in terms of classification and reproducibility for feline variants, the applicability across species was tested. Practically, five variants per species, when available, were classified and the evaluators were asked to note 1) any incompatibilities, and/or 2) difficulties not seen in the cat. For both questions, the answer was “no” for every single variant across eight additional species (i.e., dog, horse, cattle, pig, goat, sheep, rabbit, and chicken).

As there were no issues encountered, several of the remaining variants submitted for the feline pathogenic variant dataset that did not pass all criteria, were additionally evaluated, resulting in 72 feline variants or 53% (= 72/136) of the disease-associated variants currently known in cats, classified. Population data was collected for five of these additional variants, but all were observed to be homozygous wildtype. Altogether, we have now provided a classification for over 110 variants (Suppl. Table S9).

## 4 Discussion

Breeding strategies, especially in companion animals, significantly impact the health of their offspring and the overall population. Selective breeding that emphasizes specific traits too narrowly can result in reduced genetic diversity, ultimately potentially increasing the population’s susceptibility to diseases. Furthermore, prioritizing phenotypic traits over health can adversely affect animal welfare. Responsible breeding practices that emphasize health, genetic diversity, and animal welfare can lead to healthier and more resilient animal populations(74). Especially in inbred populations with a small effective population size, the use of validated genetic testing is essential. Therefore, we have developed and tested the AVCG, aiming not only to provide guidance to the practitioner who sees the individual animal in a clinical setting but also to assist in improving the health of the population, by objectifying when there is sufficient evidence for a variant to be used for screening and in breeding programs. This development process is not finished with the publication of the current set of guidelines.

Common to both ACMG and AVCG and considered good practice, guidelines are subject to change. Since the publication of the original ACMG guidelines, several of the criteria have been modified, clarified, or are considered for removal, and a thorough analysis demonstrates a considerable overlap with the decisions made here for AVCG. From the seven criteria that were removed in the process to develop the AVCG, the two linked to expert opinion (i.e., ACMG PP5 and ACMG BP6, see Suppl. Table S2) have also been recommended for removal from ACMG, with a rationale similar to ours: actual data should be preferred over opinion(19). For three others linked to AF (ACMG PM2, ACMG BS1, and ACMG BA1, see Suppl. Table S2), parallel observations have been made, but different actions have been undertaken, probably driven by the difference in occurrence. In more detail, ACMG has recognized that, for some variants, the stand-alone criterion for an AF of 5% is too strict, which has been tackled by a (very short) exception list, and it has been suggested to reduce the moderate weight of the allelic frequency criterion supporting pathogenic classification(20,21). Here, 9% (5/53) of the feline variants in the pathogenic list have AFs >5% overall or in subpopulations (with subpopulation being a breed or a variety within a breed, depending on the registry, see Suppl. Table S1 for examples) and even higher percentages were found in other population studies in cats (17%, 9/52)(33) and dogs (18%, 46/250)(75), indicating an exception list is not feasible for species or populations in which genetic diversity is reduced. For AVCG, we opted to omit the three criteria linked to AF, as explained above, and this led to a marked improved classification. Clarifications were also deemed necessary for the criteria linked to *de novo* variants (ACMG PS2 and PM6) in ACMG(22). These were also considered unclear here and led to a modification of criterion PS2 and removal of criterion PM6 (see Suppl. Table S2). As such, for six out of seven criteria that were removed, at least a modification was also deemed necessary after the original ACMG guidelines were published.

A change implemented by ACMG, but not adopted for AVCG, was the update on PVS1(23). The rationale for a more extensive explanation for this criterion for ACMG was that it is the only criterion with a very strong weight, hence it can influence the classification of a variant easily. The proposed change is an extensive flow chart, which is complex and should be re-evaluated as the AVCG evolve. Published during this study, the updated ACMG recommendations linked to all criteria associated with splice variants (PVS1, PS1, PP3, BP4, BP6) could not be included in the evaluation process(24). Nevertheless, except for small additions to PVS1 (explicitly stating that this criterion can also be used across species) and PP3/BP4 (explicitly stating the number of tools that should be used for *in silico* criteria), the group considered the criteria linked to splicing sufficiently clear. Similarly, criteria PS3/BS3, PP4/BS4, the criterion formally known as PP1 but renamed to PS5 here as the weight was increased, and PM3 were also considered clear, while additional clarifications were published for ACMG(25–27).

The final recommendation dealt with PP3 and BP4, i.e., the performance of *in silico* tools(28). Remarkably, while there are several benchmark studies, there are none for any of the species evaluated here. While we wanted to provide guidance, this posed significant challenges as there are, to our knowledge, no reference datasets for any of the animals evaluated here. To identify the most optimal strategy to develop such a dataset, systematic literature reviews were conducted. Surprisingly, the number of different strategies was limited, and especially for the benign variants, only one method remained. As the most common methodology was AF-based, with all its limitations in species with limited genetic diversity, we opted for a different strategy(6,29). This strategy has the additional benefit that it does not rely on any of the criteria used to classify variants later on, which is positive as this would have created some sort of circularity in reasoning. While this is important for the development of the AVCG, it is also a common issue when *in silico* tools are tested. Circularity occurs when there is an overlap between training and test datasets, which is common in humans, and can lead to biased performance estimates(54,76). As none of the tools used feline genetic variants during the development process, circularity is no issue here.

Overall, we observed large performance differences between and within categories of variant types (Table 6, Suppl. Table S7). Similarly, benchmarks in literature indicate even gene- and phenotype-specific difference(41–43). While individual tools might outperform one another in specific cases, we aimed to provide an overview of the combination of tools that generally demonstrated the best accuracy, which turned out to be the combination of MutPred2 and LIST-S2 for missense variants(56,62). The tools in this combination differ in underlying methodology and how they were trained, which is an advantage as this also implies less dependency on the same information, that is, a more independent analysis and conclusion. While a satisfactory performance was achieved for missense variants and splice sites, this was not the case for the other variant types. As information from variant effect predictor tools can be readily generated, we, in general, encourage the use of these kind of tools if performance is adequate, which is thus limited to missense variants and splice sites based on the current benchmark.

Combined, classification of animal variants with AVCG outperformed classification with ACMG. When implemented in practice, however, guidelines ideally also have an additional characteristic, i.e., a high reproducibility or concordance between evaluators. If this is not the case, results tend to change between laboratories or even between evaluators within a laboratory. While interpretation differences have been reported several times, variant classification reproducibility has been demonstrated to be rather low with and without the ACMG guidelines (34% for both)(3–5). A cautionary note on this result is that a disagreement of 66% does not necessarily imply also that clinical management will be affected in two out of three cases. Looking at differences that might affect clinical management (L/LP versus VUS/LB/B), the number reduced to 22%. The difference with AVCG is remarkable: an exact agreement of 76% and only 4% of the differences might have practical consequences. While an obvious explanation might be that only variants that were easy to classify were included, this was not the case. In fact, even when the comparison solely includes the group of variants among which there was debate, the agreement was still twice as high with the AVCG (ACMG: 34% versus AVCG: 60%).

With this study, we not only provide methods to develop datasets for benchmarking, but we also classify and provide AFs, and *in silico* data for over half of the currently known feline pathogenic variants. Overall, nearly all variants classified here got an LP or P label, and for several of the ones that did not (e.g., three hypertrophic cardiomyopathy-associated variants), this was anticipated(6,29,77,78). The high number of LPs/Ps was expected as the study was based on OMIA, a database that focuses on disease-associated variants. The range of phenotypes, inheritance patterns, penetrance, and species in which the AVCG were successfully applied, indicates that the central scope, i.e., providing guidelines that can be used in general for Mendelian disorders, seems to be fulfilled. However, the classification of these variants is not written in stone. Similarly to the guidelines themselves and the results obtained with the *in silico* tools, the classification of a variant can change when new information becomes available(33). One example is the UROS:c.140C>T variant. In the original publication that was also used to classify the variant here, one cat was homozygous for two UROS variants (UROS:c.140C>T and UROS:c.331G>T)(68). As functional studies indicated a potential effect for both variants (with a more pronounced effect for the latter), the effect of the two could not be separated(68). Recently, several cats without symptoms but homozygous for the UROS:c.140C>T variant were identified, indicating a re-evaluation needs to be done(33). This substantiates the importance of follow-up, but also the importance of identifying additional cases, whenever possible.

This study raises interesting questions for future research. As some disease-causing variants were classified as VUS with AVCG, further optimization and tailoring, especially for the n=1 situation, should be a future focus. Furthermore, while a large set of variants was thoroughly checked, the aim should be to try to classify all variants currently published. This will undoubtedly identify additional areas of improvement and potential classification difficulties. Importantly, while striving for perfect accuracy in terms of classification, we also want to stress that there will likely always be exceptions that do not follow the rules. Finally, we agree with the view of ACMG and that is why these guidelines focus specifically on Mendelian diseases and why variants associated with complex diseases, somatic variants, and structural variants larger than 50bp, were excluded(2). While the excluded variants currently represent a minority of the total number of disease-associated variants, more are likely to be discovered in the future, requiring solutions for classification for them as well. As such, we also support the foundation of expert groups, an initiative which is currently underway under the umbrella of the International Society for Animal Genetics, to take on the challenge of 1) further improving guidelines, 2) keeping track of new data and, whenever necessary, updating variant classification, and 3) developing guidelines for the variants that were outside the current scope. Furthermore, steps are undertaken to make these variant classifications publicly available in databases like OMIA.

In short, we provide the AVCG, tailored for variant classification in animals, and demonstrate a substantially improved classification, as well as reproducibility, even when used on animal variants the ACMG guidelines struggle with or on animal variants which led to individual conflicting assessments.

## 6 Author Contributions

BJGB conceived and designed the analysis. FB, MA, HA, IC, CDdC, JJH, JH, MDK, EL, IL, ML, LAL, ÅO, LP, PS, TV, FGvS, BJGB provided and/or reviewed putative pathogenic variants. FB, MA, HA, JJH, JH, MDK, EL, IL, ML, LAL, ÅO, LP, PS, TV, FGvS, BJGB participated in the development process of the guidelines. HA and CDdC collected the allele frequency data. FB, MA, IC, JJH, ÅO, FGvS and BJGB performed the classification process. FB, MA, IC, JJH, FGvS and BJGB executed the reproducibility study. FGvS and BJGB wrote the original draft of the paper. All authors reviewed, improved and approved the manuscript.

## Acknowledgments

We would like to thank the Bijzonder Onderzoeksfonds (BOF) for providing the funding for the PhD grant of the first author.

## 8 Data Availability Statement

All relevant data is contained within the article: The original contributions presented in the study are included in the article/supplementary material, further inquiries can be directed to the corresponding author.

