## Supplementary Table S2 for "Variant classification guidelines for animals to objectively evaluate genetic variant pathogenicity"

Suppl. Table S2. Overview of the original American College of Medical Genetics and Genomics (ACMG) guidelines and summary of changes.

| ACMG name | AVCG name | Criterion |
| --- | --- | --- |
| PVS1 | **PVS1** | Null variant (nonsense, frameshift, canonical ±1 or 2 splice sites, initiation codon, single or multiexon deletion) in a gene where LOF is a known mechanism of disease. |
| PS1 | PS1 | Same amino acid change as a previously established pathogenic variant regardless of nucleotide change. |
| PS2 | **PS2** | *De novo* (both maternity and paternity confirmed) in a patient with the disease and no family history. |
| PS3 | PS3 | Well-established *in vitro* or *in vivo* functional studies supportive of a damaging effect on the gene or gene product. |
| PS4 | PS4 | The prevalence of the variant in affected individuals is significantly increased compared with the prevalence in controls. |
| PM1 | **PM1** | Located in a mutational hot-spot and/or critical and well-established functional domain (e.g., active site of an enzyme) without benign variation. |
| PM2 | NA | Absent from controls (or at extremely low frequency if recessive) (Table 6) in Exome Sequencing Project, 1000 Genomes Project, or Exome Aggregation Consortium. |
| PM3 | PM3 | For recessive disorders, detected in *trans* with a pathogenic variant. |
| PM4 | PM4 | Protein length changes as a result of in-frame deletions/insertions in a nonrepeat region or stop-loss variants. |
| PM5 | **PM2** | Novel missense change at an amino acid residue where a different missense change determined to be pathogenic has been seen before. |
| PM6 | NA | Assumed *de novo*, but without confirmation of paternity and maternity. |
| PP1 | **PS5** | Cosegregation with disease in multiple affected family members in a gene definitively known to cause the disease. |
| PP2 | PP2 | Missense variant in a gene that has a low rate of benign missense variation and in which missense variants are a common mechanism of disease. |
| PP3 | **PP3** | Multiple lines of computational evidence support a deleterious effect on the gene or gene product (conservation, evolutionary, splicing impact, etc.) |
| PP4 | PP4 | Patient’s phenotype or family history is highly specific for a disease with a single genetic etiology. |
| PP5 | NA | Reputable source recently reports variant as pathogenic, but the evidence is not available to the laboratory to perform an independent evaluation. |
| BA1 | NA | Allele frequency is >5% in Exome Sequencing Project, 1000 Genomes Project, or Exome Aggregation Consortium. |
| BS1 | NA | Allele frequency is greater than expected for disorder. |
| BS2 | BS2 | Observed in a healthy adult individual for a recessive (homozygous), dominant (heterozygous), or X-linked (hemizygous) disorder, with full penetrance expected at an early age. |
| BS3 | BS3 | Well-established *in vitro* or *in vivo* functional studies show no damaging effect on protein function or splicing. |
| BS4 | BS1 | Lack of segregation in affected members of a family. |
| BP1 | NA | Missense variant in a gene for which primarily truncating variants are known to cause disease. |
| BP2 | BP2 | Observed in *trans* with a pathogenic variant for a fully penetrant dominant gene/disorder or observed in *cis* with a pathogenic variant in any inheritance pattern. |
| BP3 | BP3 | In-frame deletions/insertions in a repetitive region without a known function. |
| BP4 | **BP4** | Multiple lines of computational evidence suggest no impact on gene or gene product (conservation, evolutionary, splicing impact, etc.). |
| BP5 | BP5 | Variant found in a case with an alternate molecular basis for disease. |
| BP6 | NA | Reputable source recently reports variant as benign, but the evidence is not available to the laboratory to perform an independent evaluation. |
| BP7 | BP6 | A synonymous (silent) variant for which splicing prediction algorithms predict no impact to the splice consensus sequence nor the creation of a new splice site AND the nucleotide is not highly conserved. |

In the first column, the original ACMG name is reported. The second column depicts whether the criterion was removed (NA), altered (**AVCG criterion name in in bold**) or remained unchanged (AVCG criterion name in normal typeset).
