## Supplementary Table S7 for "Variant classification guidelines for animals to objectively evaluate genetic variant pathogenicity"

Suppl. Table S7. Pairwise combinations of *in silico* tools that were able to score all variants in their category.

| Tool 1 | Tool 2 | Type | N_benign_ correct | N_patho_ correct | N_total_ correct | Incorrect | Inconsist | Accuracy (%) |
| --- | --- | --- | --- | --- | --- | --- | --- | --- |
| PhD-SNP | SNPs&Go | missense | 19 | 13 | 32 | 4 | 6 | 32/36 (89) |
| PhD-SNP | MutPred2 | missense | 19 | 16 | 35 | 3 | 4 | 35/38 (92) |
| PhD-SNP | PredictSNP | missense | 18 | 16 | 34 | 4 | 4 | 34/38 (89) |
| PhD-SNP | LIST-S2 | missense | 18 | 14 | 32 | 1 | 9 | 32/33 (97) |
| PhD-SNP | MetaSNP | missense | 18 | 15 | 33 | 4 | 5 | 33/37 (89) |
| SNPs&Go | MutPred2 | missense | 21 | 13 | 34 | 4 | 4 | 34/38 (89) |
| SNPs&Go | PredictSNP | missense | 19 | 13 | 32 | 4 | 6 | 32/36 (89) |
| SNPs&Go | LIST-S2 | missense | 20 | 10 | 30 | 1 | 11 | 30/31 (97) |
| SNPs&Go | MetaSNP | missense | 19 | 13 | 32 | 5 | 5 | 32/37 (86) |
| MutPred2 | PredictSNP | missense | 19 | 17 | 36 | 4 | 2 | 36/40 (90) |
| **MutPred2** | **LIST-S2** | **missense** | **20** | **14** | **34** | **1** | **7** | **34/35 (97)** |
| MutPred2 | MetaSNP | missense | 19 | 16 | 35 | 4 | 3 | 35/39 (90) |
| PredictSNP | LIST-S2 | missense | 18 | 14 | 32 | 1 | 9 | 32/33 ( 97) |
| PredictSNP | MetaSNP | missense | 19 | 16 | 35 | 6 | 1 | 35/41 (85) |
| LIST-S2 | MetaSNP | missense | 18 | 13 | 31 | 1 | 9 | 31/32 (97) |
| **Spliceator** | **SSPnn** | **splice sites** | **NA** | **NA** | **6** | **0** | **2** | **6/6 (100)** |
| Spliceator | GENSCAN | splice sites | NA | NA | 3 | 0 | 5 | 3/3 (100) |
| SSPnn | GENSCAN | splice sites | NA | NA | 5 | 0 | 3 | 5/5 (100) |

N_benign_ correct = number of neutral variants correctly assessed, N_patho_ correct = number of pathogenic variants correctly assessed; N_total_ correct = total number of variants correctly assessed (sum of N_benign_ correct and N_patho_ correct); Incorrect = number of variants consistently classified wrong; Inconsist =number of variants for which the result of both tools conflicted; Accuracy = N_total_ correct/ (N_total_ correct + Incorrect). As some variants remove splice (acceptor or donor) sites and others add splice sites, “benign” and “patho” does not accurately reflect the action, hence, the total number of variants correctly assessed, is provided for the splice site tools. The combinations that scored best for the two types of variants, are depicted in bold.
