## Supplementary Table S8 for "Variant classification guidelines for animals to objectively evaluate genetic variant pathogenicity"

Suppl. Table S8. Variants used to quantify reproducibility.

| Variant | Effect | Reason for inclusion | Classification | *In silico* criterion |
| --- | --- | --- | --- | --- |
| MAN2B1:c.1749_1752delCCAG | Frameshift | Confident pathogenic | P – P – P | See Suppl. Table S2 |
| CYB5R3:g.137967506CT | Missense | Confident pathogenic | LP – L P – P | See Suppl. Table S2 |
| F9:g.117111577CT | Nonsense | Confident pathogenic | P – P – P | See Suppl. Table S2 |
| HEXB:g.141571030_141571044del | Splice site | Confident pathogenic | P – P – P | See Suppl. Table S2 |
| HEXB:g.141571788_141571812inv | Frameshift | Confident pathogenic | P – P – P | See Suppl. Table S2 |
| CYB5R3:g.137970815CG | Splice site | Confident pathogenic | P – P – P | See Suppl. Table S2 |
| MYBPC3:c.91G > C | Missense | Difficult to classify with ACMG | P – P – P | Disease |
| MYBPC3:c.220G > A | Missense | Difficult to classify with ACMG | VUS – VUS – LB | Benign |
| MYBPC3:c.2453C > T | Missense | Difficult to classify with ACMG | P – P – P | Disease |
| TNNT2:c.95-108G > A | Splice site | Difficult to classify with ACMG | VUS – VUS - VUS | Inconsistent |
| ALMS1:c.7384G > C | Missense | Difficult to classify with ACMG | VUS – VUS - VUS | Disease |
| MYH7:c.5647G > A | Missense | Difficult to classify with ACMG | P – LP – LP | Disease |
| **CYP27B1:g.86180281del** | **Frameshift** | **Disagreement among evaluators** | **VUS – VUS – P** | NA |
| COL5A1:g.93210344del | Frameshift | Disagreement among evaluators | P – LP – P | NA |
| CREB3L1:g.100436508_100436509del | Frameshift | Disagreement among evaluators | P – P – LP | NA |
| ATP7B :g.19611002C>G | Missense | Disagreement among evaluators | LP – LP – LP | Disease |
| TAC3:g.85517451C>T | Missense | Disagreement among evaluators | VUS – VUS - VUS | Benign |

Seventeen variants were included, of which five were frameshifts, one was a nonsense, eight were missense and three were splice sites. The label assigned with the Animal Variant Classification Guidelines (AVCG) by each evaluator is provided (“Classification”), as well as the reason to include the variant in the reproducibility analysis (“Reason for inclusion”). When applicable (i.e., for missense and splice site variants) and when not already mentioned in Suppl. Table S2, the outcome of the combination of *in silico* variant effect predictor tools is also reported (“*in silico* criterion”; for missense: based on combination of LIST-S2 and MutPred2; for splice sites: based on combination of Spliceator and SSPnn). Variants in which the disagreement might lead to changes in clinical management, screening, and/or breeding, are marked in bold. Abbreviations: ACMG = American College of Medical Genetics and Genomics, P = pathogenic, LP = likely pathogenic, VUS = variant of unknown significance, B = benign, LB = likely benign, NA = not applicable.
