## Supplementary Data S1 - S5 and Supplementary Figures S1-S2 for "Variant classification guidelines for animals to objectively evaluate genetic variant pathogenicity"

Supplementary Material

### 1 Supplementary Data

Suppl. Data S1. Overview of the systematic review strategy to identify pathogenic and benign variants, according to the PRISMA guidelines.

- Type of review: systematic review
- Primary research question: Which strategies are used in benchmark studies to identify pathogenic and benign/neutral variants?
- Search strategy:
  - 1. Database: PubMed
  - 2. Websites: https://www.varianteffect.org/veps
  - Inclusion criteria: all articles that are included should describe a selection strategy for pathogenic or benign variants to assess performance of a variant effect predictor
  - Exclusion criteria:
    - No mention of a selection strategy for benign and/or neutral variants
- Query string:

("missense"[Title/Abstract] OR "nonsense"[Title/Abstract] OR "in-frame"[Title/Abstract] OR "frameshift"[Title/Abstract] OR "splice site"[Title/Abstract]) AND ("benchmark" OR "performance") AND ("variant effect predictor"[Title/Abstract] OR "in silico"[Title/Abstract]) NOT ("cancer*" OR "tumor" OR "tumour") and date set to prior 2024-01

- Search validation procedure (check for bias):

The following reports should be retrieved:

1. Ghosh R, Oak N, Plon SE. Evaluation of in silico algorithms for use with ACMG/AMP clinical variant interpretation guidelines. Genome Biol. 2017 Nov 28;18(1):225. doi: 10.1186/s13059-017-1353-5. PMID: 29179779; PMCID: PMC5704597.
2. Thusberg J, Olatubosun A, Vihinen M. Performance of mutation pathogenicity prediction methods on missense variants. Hum Mutat. 2011 Apr;32(4):358-68. doi: 10.1002/humu.21445. Epub 2011 Feb 22. PMID: 21412949.

All aforementioned reports were found.

Suppl. Data S2. Overview of the systematic review strategy to identify *in silico* tools, according to the PRISMA guidelines.

- Type of review: systematic review
- Primary research question:
  - Which variant effect predictors exist?
  - Which ones have an online user interface?
  - Which of the remaining ones can be used for animals?
- Search strategy:
  - 1. Database: PubMed
  - 2. Websites: https://www.varianteffect.org/veps
  - Inclusion criteria: all articles that are included should describe a tool that predicts the functional impact and accepts cat proteins, which is easily accessible (i.e., a web-tool where you can submit the variant immediately without a download)
  - Exclusion criteria:
    - No web-tool
    - Cannot be used for animals
    - Tool requiring a fee
- Query string:

("missense"[Title/Abstract] OR "nonsense"[Title/Abstract] OR "in-frame"[Title/Abstract] OR "frameshift"[Title/Abstract] OR "splice site"[Title/Abstract]) AND ("benchmark" OR "performance") AND ("variant effect predictor"[Title/Abstract] OR "in silico"[Title/Abstract]) NOT ("cancer*" OR "tumor" OR "tumour") and date set to prior 2024-01

- Search validation procedure (check for bias):

All tools reported in the veterinary literature should be retrieved based on these results. These were MaxEntScan, NNSplice, PolyPhen-2, SNAP, PROVEAN, SIFT, MutPred, PredictSNP, CADD, Meta-SNP and Condel. All tools were found.

Suppl. Data S3. Explanatory comments of criteria in the Variant Classification Guidelines.

**PVS1**
The group finds it important to specify that evidence for loss-of-function as a known mechanism can be based on the same or other species, but if it is across species, it is also important to evaluate whether it is likely that the function of the gene is similar. The reason to include this “function” part is e.g., that some genes are pseudogenes in one species and not in another, something which is more likely to occur if species are distantly related. If a gene is expected, or has been shown to be functional in the species studied, this criterion can be used. From the description, it is clear that this criterion applies to variants that disrupt gene function. This is not always easy to assess: e.g., the presence of alternative transcripts can complicate interpretation and some transcripts are tissue-specific.

**PS1**
If a variant results in an amino-acid change that has already been shown to be pathogenic, this criterion applies. If a variant results in a **different** amino-acid than the one that has been shown to be pathogenic, it is not this criterion that applies, but PM2.

**PS2**
It is important to evaluate this vertically throughout the pedigree: parents should test negative for the variant. If the parents are not available or were not tested, this criterion cannot be used.

**PS3**
No additional explanation.

**PS4**
The group finds it difficult to confidently decide on which odds ratio (OR) is sufficiently high and at the same time to be of practical use as a cut-off. Deciding on which cut-off is appropriate has a lot of consequences: large sample sizes can lead to significant results with relatively low point estimates and vice versa. In human studies, an OR of 5 has been suggested to be an appropriate cut-off. Studies in animals suggest a similar cut-off would also be appropriate. An explanation on how to calculate the various ORs is provided (Suppl. Data S5*)*. There are two important remarks for this criterion:

- this criterion cannot be used for *de novo* variants as there is no true “enrichment” in cases: it typically will only occur once. For *de novo* variants, the PS2 criterion should thus be used.
- it is important to ensure an adequate sample size is available. A power analysis should be performed.

**PS5**
Practically, this implies that in one or more pedigrees, the variant is present in a disease-causing state in affected family members and that the variant is not present in a disease-causing state in healthy individuals. When evaluating this criterion, it is important to take into account that the litter size can vary across species and even across breeds within a species as this influences the size of the pedigree that can reasonably be expected.

**PM1**
No additional explanation.

**PM2**
The group finds it important to specify that this also involves across species.

**PM3**
Several remarks are important in this criterion. Firstly, both variants should be reported jointly to allow an assessment of the association. Secondly, it may be difficult to assess the effect of each individual variant: some only jointly have an effect. Thirdly, sometimes phenotypes, even in the same gene, can be inherited in an autosomal dominant or an autosomal recessive manner, depending on the variant. Practically, this means that an additional variant, next to a pathogenic variant that turns out to cause a phenotype in a dominant manner, might erroneously be labelled "pathogenic". The final remark is that, although this might be a valid criterion, it should not be weighted too high for final classification of a variant.

**PM4**
It is important to emphasize that this criterion does **not** apply for variants in non-conserved regions.

**PP1/BP1**
The criterion was actually worded as “*Cross-species alignment to determine whether a variant is conserved (= supportive for pathogenic) or not conserved (=supportive for benign).* This criterion can support benign or pathogenic classification depending on the answer. It does not refer to *in silico* tools that predict the effect of a variant, because that falls under PP3 or BP4. It does refer to 1/ a cross-species sequence alignment to evaluate conservation of nucleotides or amino-acids (depending on what is relevant for that specific variant) **and** 2/ a cross-species check of the potential role of the variant based on other information, like ClinVar (e.g., if the variant has been described in ClinVar, is it considered a pathogenic or benign variant there).

**PP2**
The group finds it important to specify that this also involves across species.

**PP3/BP4**

Multiple **different** software tools support prediction. While it is difficult to objectively quantify what the optimal number of tools is as this also depends on how similar the tools are and their availability, the group decided that at least two different tools, if available, have to be used. Only if **all** tools lead to the same conclusion (i.e., when two are used, both say deleterious or both say harmless), the criterion is accordingly fulfilled. If tools are conflicting, the criterion is not fulfilled. An overview of the tools that support the use of non-human data per variant type is provided in Table 6. Furthermore, we recommend checking benchmark studies to select the most accurate tools and to keep in mind that for some variant types, *in silico* tools have a poor accuracy.

**PP4**
This criterion only applies when (i) most patients test positive for a pathogenic variant in that gene; (ii) the patient has a well-defined syndrome with little overlap with other clinical presentations. One example is Duchenne muscular dystrophy, in which a variant in the *DMD* gene is expected. Only very limited genetic heterogeneity can thus be present for this criterion to be valid. When one observes the phenotype, one has immediately one, at most few, gene(s) in mind.

**BS1**
Caveat: the presence of phenocopies for common phenotypes (e.g., epilepsy) can mimic lack of segregation among affected individuals.

**BS2/BS3**
No additional explanation.

**BP1**
see PP1.

**BP2/BP3/BP5/BP6**

No additional explanation.

Suppl. Data S4. Reporting recommendations and quality checks when publishing disease-associated variants.

In what follows, we provide some general recommendations, based on difficulties repeatedly encountered during the review process.

**Statistical set-up and calculations**

To describe genotype-phenotype associations, usually odds ratios and relative risks are used. To allow a check of these data, a 2x3 table describing the distribution of genotypes across the phenotypical categories is important. Furthermore, as various methods to perform the calculations exist, we recommend a clear description of what is exactly calculated. For odds ratios, we provide examples and terminology in Suppl. Data S4.

**Variant description**

Variant descriptions are often incomplete and do not contain version numbers. As new annotations and reference genomes are constantly published, it can be difficult to decipher exactly which variant is described. Aside from descriptions according to the accepted nomenclature, we in addition suggest to provide the neighbouring amino acids and/or nucleotides (what is most appropriate) as well as this facilitates identification of the important variant. This can for example be done in a figure describing cross-species alignment and/or electropherograms.

**Mode of inheritance, disease penetrance and genetic heterogeneity**

The distribution of genotypes across the phenotypical categories should allow a substantiated proposal for a mode of inheritance and furthermore detail whether penetrance (defined as the probability to identify the phenotype, given a genotype) is expected to be complete or reduced, as well as whether genetic heterogeneity is expected. These three items are important during the evaluation of several criteria and should thus be mentioned in the manuscript.

**Population study**

While allele frequency has been removed from the AVCG as criterion influencing classification, all group members agreed that population studies are of paramount importance when new variants are reported or already published variants investigated further. Population studies provide an overview of how widespread a variant is, i.e., whether it is case-, family-, breed-specific or common overall. Furthermore, when breeding strategies are developed, the prevalence of a disease-causing variant is important to decide on the strategy.

Suppl. Data S5. Clarification of methods to calculate odds ratios.

|  | Genotypes | | |
| --- | --- | --- | --- |
|  | Vt/Vt | Wt/Vt | Wt/Wt |
| Cases | a | b | c |
| Controls | d | e | f |

With a – f representing the number of individuals in each cell.

Several methods exist to calculate odds ratios (ORs). We specify the following two:

1/ allelic OR: this requires calculation of allelic frequencies and hence collapsing of the table

|  | Allele | |
| --- | --- | --- |
|  | Vt | Wt |
| Cases | 2 x a + b = g | 2 x c + b = h |
| Controls | 2 x d + e = i | 2 x f + e = j |

Based on this table, the OR is calculated as:

$$OR =\frac{(g \times j)}{(i \times h)}$$

2/ genotype OR: this requires a specification of a mode of inheritance and collapsing of the genotypes accordingly. In more detail,

- for an autosomal recessive mode of inheritance, assuming Vt/Vt is affected:

|  | Genotype | |
| --- | --- | --- |
|  | Vt/Vt | Wt/Wt + Wt/Vt |
| Cases | a | b + c = h |
| Controls | d | e + f = j |

Based on this table, the OR is calculated as:

$$OR =\frac{(a \times j)}{(d \times h)}$$

- for an autosomal dominant mode of inheritance, assuming Wt/Vt and Vt/Vt are affected:

|  | Genotype | |
| --- | --- | --- |
|  | Wt/Vt + Vt/Vt | Wt/Wt |
| Cases | a + b = g | c |
| Controls | d + e = i | f |

Based on this table, the OR is calculated as:

$$OR =\frac{(g \times f)}{(i \times c)}$$

Remark:

When one of the cells is zero, the OR becomes zero as well or is not defined. One can apply at that moment a correction by adding 0.5 to every cell. In what follows, we provide an example, based on the WNK4:c.2899C>T variant, which follows an autosomal recessive mode of inheritance, with the following distribution of genotypes over cases and controls:

|  | Genotypes | | |
| --- | --- | --- | --- |
|  | Vt/Vt | Wt/Vt | Wt/Wt |
| Cases | 43 | 0 | 0 |
| Controls | 0 | 22 | 69 |

- For the allelic OR, the table collapses to:

|  | Allele | |
| --- | --- | --- |
|  | Vt | Wt |
| Cases | 43 x 2 = 86 | 0 |
| Controls | 22 | 69 x 2 + 22 = 160 |

As the OR is undefined in this example, we add 0.5 to every cell count:

|  | Allele | |
| --- | --- | --- |
|  | Vt | Wt |
| Cases | 86.5 | 0.5 |
| Controls | 22.5 | 160.5 |

The allelic OR now has a value of 1,234.067

- For an autosomal recessive mode of inheritance, the genotype OR table has the following values:

|  | Genotype | |
| --- | --- | --- |
|  | Vt/Vt | Wt/Wt + Wt/Vt |
| Cases | 43 | 0 |
| Controls | 0 | 91 |

As the OR is undefined in this example, we add 0.5 to every cell count:

|  | Genotype | |
| --- | --- | --- |
|  | Vt/Vt | Wt/Wt + Wt/Vt |
| Cases | 43.5 | 0.5 |
| Controls | 0.5 | 91.5 |

The genotype OR now has a value of 15,921.

Suppl. Figure S1. PRISMA 2020 flow diagram for new systematic reviews which included searches of databases, registers and other sources for selection strategies of benign and pathogenic variants.

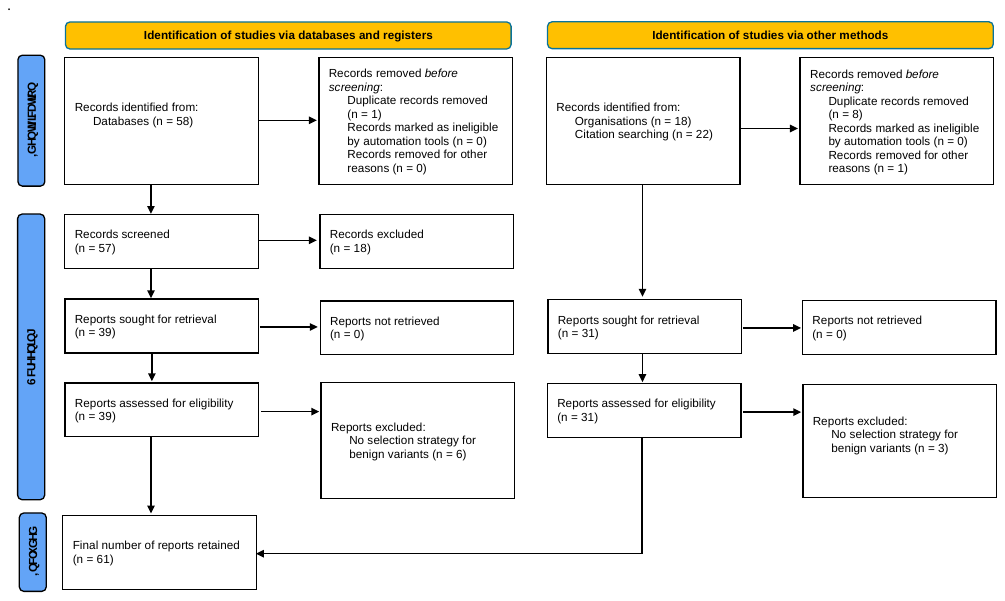

Suppl. Figure S2. PRISMA 2020 flow diagram for new systematic reviews which included searches of databases, registers and other sources for selection strategies of *in silico* tools.

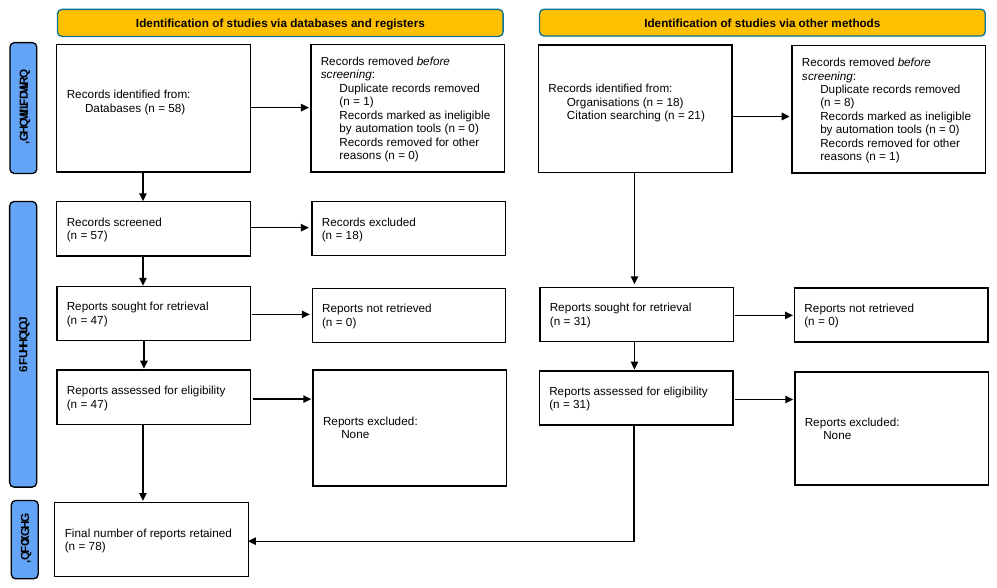
